# Synergistic cues from diverse bacteria enhance multicellular development in a choanoflagellate

**DOI:** 10.1101/851824

**Authors:** Ella V. Ireland, Arielle Woznica, Nicole King

## Abstract

Bacteria regulate the life histories of diverse eukaryotes, but relatively little is known about how eukaryotes interpret and respond to multiple bacterial cues encountered simultaneously. To explore how a eukaryote might respond to a combination of bioactive molecules from multiple bacteria, we treated the choanoflagellate *Salpingoeca rosetta* with two sets of bacterial cues, one that induces mating and the other that induces multicellular development. We found that simultaneous exposure to both sets of cues enhanced multicellular development in *S. rosetta*, eliciting both larger multicellular colonies and an increase in the number of colonies. Thus, rather than conveying conflicting sets of information, these distinct bacterial cues synergize to augment multicellular development. This study demonstrates how a eukaryote can integrate and modulate its response to cues from diverse bacteria, underscoring the potential impact of complex microbial communities on eukaryotic life histories.

**Importance:** Eukaryotic biology is profoundly influenced by interactions with diverse environmental and host-associated bacteria. However, it is not well understood how eukaryotes interpret multiple bacterial cues encountered simultaneously. This question has been challenging to address because of the complexity of many eukaryotic model systems and their associated bacterial communities. Here, we studied a close relative of animals, the choanoflagellate *Salpingoeca rosetta*, to explore how eukaryotes respond to diverse bacterial cues. We found that a bacterial chondroitinase that induces mating on its own can also synergize with bacterial lipids that induce multicellular “rosette” development. When encountered together, these cues enhance rosette development, resulting in the formation of more rosettes, each containing more cells than rosettes that form in the absence of the chondroitinase. These findings highlight how synergistic interactions among bacterial cues can influence the biology of eukaryotes.

## Introduction

Eukaryotes, including animals and their closest living relatives, choanoflagellates, encounter abundant and diverse bacteria in the environment (1–3). However, interactions among eukaryotes and bacteria can be challenging to study in animal models due to the complex physiology of the hosts and the large number of oftentimes unculturable bacteria present, each of which releases diverse molecules (4–6). Multiple types of intestinal bacteria are required to induce full immune maturation in mice and humans, but it remains unclear whether this is due to interactions among the bacteria, or the integration by the host of multiple independent bacterial cues (7–11). The interaction of a eukaryote with multiple partners can change the magnitude or directionality of each pair-wise interaction (12), and it can be challenging to measure the functional and fitness effects of such complex networks (13). Therefore, simpler model systems may be necessary to investigate how animals and other eukaryotes integrate information from multiple bacterial cues encountered at the same time.

The choanoflagellate *Salpingoeca rosetta* can serve as a simple model for studying interactions between bacteria and eukaryotes. Like all choanoflagellates, *S. rosetta* captures bacterial prey from the water column using an apical “collar complex” composed of a microvillar collar surrounding a single flagellum (Fig. 1A; (14, 15)). In addition, like many animals (2, 16, 17), *S. rosetta* undergoes important life history transitions in response to distinct bacterial cues. For example, a secreted bacterial chondroitinase called EroS (for <u>E</u>xtracellular <u>R</u>egulator <u>o</u>f <u>S</u>ex) produced by *Vibrio fischeri*, *Proteus vulgaris*, and select other Gammaproteobacteria induces solitary *S. rosetta* cells to gather into mating swarms (Fig. 1B; (18)). The cells in mating swarms are not stably adherent and eventually resolve into pairs of cells that mate by undergoing cell and nuclear fusion, followed by meiotic recombination. When exposed to a different type of bacterial cue, specific sulfonolipids called RIFs (for <u>R</u>osette Inducing <u>F</u>actors) from the Bacteroidetes bacterium *Algoriphagus machipongonensis*, solitary cells of *S. rosetta* undergo serial rounds of cell division without separation, thereby resulting in the development of multicellular rosettes of cells (Fig. 1C) that are physically linked by cytoplasmic bridges and a shared extracellular matrix (19–22).

**Figure 1:**
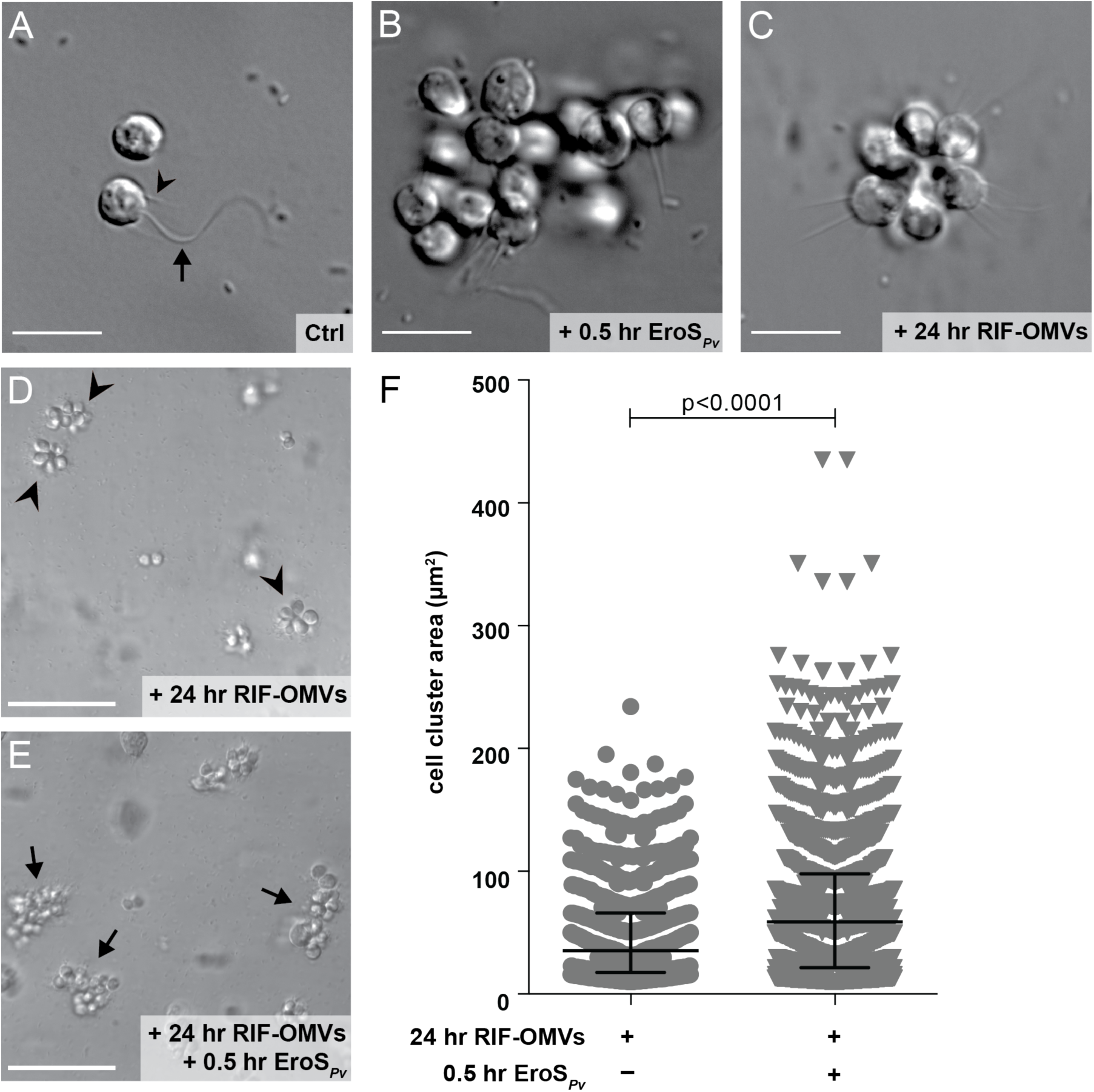
Rosettes swarm in response to the EroS_*Pv*_ mating factor. (A-C) Bacterial cues regulate mating and multicellularity in *S. rosetta*. Scale bars = 10 µm. (A) *S. rosetta* grown in the presence of the prey bacterium *E. pacifica* (“Ctrl”) proliferated as solitary cells. This culture served as the foundation for all experiments in this study. A typical *S. rosetta* cell has an apical collar (arrowhead) surrounding a single flagellum (arrow). (B) *S. rosetta* formed mating swarms within 0.5 hours of treatment with the bacterially-produced chondroitinase EroS_*Pv*_. (C) *S. rosetta* solitary cells developed into rosettes through serial rounds of cell division within 24 hours of treatment with RIF-OMVs from the bacterium *A. machipongonensis*. (D-E) Rosettes swarm in the presence, but not in the absence, of EroS_*Pv*_. Scale bars = 50 µm. (D) After 24 hours of treatment with RIF-OMVs, solitary cells in an SrEpac culture developed into rosettes (arrowheads) but did not swarm. (E) Swarms of rosettes (arrows) formed after 24 hours of treatment with RIF-OMVs followed by 0.5 hours of treatment with EroS_*Pv*_. (F) Shown are the surface areas of cell clusters from SrEpac cultures treated with RIF-OMVs for 24 hours followed by 0.5 hours of incubation either with or without EroS_*Pv*_. Following the approach of (18), we generated a binary mask to measure cell cluster area (the area of each cell, rosette, or swarm; Fig. S1). EroS_*Pv*_ treatment resulted in clusters of cells, including swarms of rosettes (median = 58.7 µm^2^, interquartile range = 21.6-98.0 µm^2^), whose areas were significantly larger than those measured in the rosette-only control (median = 35.5 µm^2^, interquartile range = 17.8-65.9 µm^2^) (Kolmogorov-Smirnov test). 875 cell cluster areas were plotted for the cultures treated with RIF-OMVs and 1359 cell cluster areas were plotted for the cultures treated with RIF-OMVs + EroS_*Pv*_.

Mating and rosette development in *S. rosetta* differ in many respects, including the chemical nature of the bacterial cues (a protein versus lipids) and the underlying cell biology (cell aggregation versus incomplete cytokinesis). Moreover, the time scales of these processes differ, with mating swarms forming within 0.5 hours of EroS treatment (18), while definitive rosettes require multiple rounds of cell division and are not observed until 11 - 24 hours after exposure to RIFs (19–22).

Motivated by the existence of distinct *S. rosetta* life history transitions that can be regulated by biochemically unrelated bacterial cues, we used *S. rosetta* as a simple model for exploring how eukaryotes are influenced by environments filled with diverse bacterial cues. We investigated how *S. rosetta* responds to environments containing both the mating inducer EroS and the rosette-inducing RIFs. We found that the initiation of mating behavior is unchanged in the presence of cues that induce rosette development. In contrast, rosette development is significantly enhanced by the presence of the mating inducer, revealing that *S. rosetta* integrates information from seemingly unrelated bacterial cues during rosette development.

## Results

### Rosettes swarm in response to the EroS_*Pv*_ mating factor

In a culture containing *S. rosetta* and the prey bacterium *Echinicola pacifica* (together comprising a culture called SrEpac; (23, 24)), solitary cells proliferated rapidly, but underwent no other observable cell state transitions (Fig. 1A). When the SrEpac culture was treated with the secreted bacterial chondroitinase EroS from *P. vulgaris* (EroS_*Pv*_)*, S. rosetta* cells formed mating swarms of 2-50 cells within 0.5 hours (Fig. 1B, Table 1), as previously reported (18). In contrast, treatment of SrEpac with *A. machipongonensis* RIFs contained in <u>o</u>uter <u>m</u>embrane <u>v</u>esicles (RIF-OMVs) induced development of multicellular rosettes within 24 hours (Fig. 1C, D, Table 1; (19, 22)).

**Table 1:** *S. rosetta* phenotypes induced by EroS_*Pv*_ and RIF-OMVs

| **Bacterial cue** | **Hours after induction** | ***S. rosetta* phenotype** | **Effect on swarming** | **Effect on rosette development** |
| --- | --- | --- | --- | --- |
| EroS<sub>PV</sub> | 0.5 | swarming | induces | n/a |
| RIF-OMVs | 0.5 | solitary | none | n/a |
| EroS<sub>PV</sub> + RIF-OMVs | 0.5 | swarming | induces | n/a |
| EroS<sub>PV</sub> | 24 | swarming | induces | none |
| RIF-OMVs | 24 | rosette | none | induces |
| EroS<sub>PV</sub> + RIF-OMVs | 24 | rosette + swarming | induces | enhances |

We then tested how mature rosettes (formed in response to pre-treatment with RIF-OMVs for 24 hours) would respond to the mating inducer EroS. After treatment with EroS_*Pv*_ for 0.5 hours, the pre-formed rosettes gathered into swarms that were quantifiable by their increase in area (median = 58.7 µm^2^, interquartile range = 21.6-98.0 µm^2^) as compared to untreated rosettes (median = 35.5 µm^2^, interquartile range = 17.8-65.9 µm^2^) (Fig. 1D-F, Table 1). Therefore, rather than being mutually exclusive, the rosette morphology induced by RIF-OMVs and the swarming behavior induced by EroS_*Pv*_ are compatible. This indicates that cells in a life history stage induced by one bacterial cue (in this case RIF-OMVs) can respond to a second bacterial cue (EroS_*Pv*_). Swarms of choanoflagellate rosettes have not previously been reported, to our knowledge, and their ecological relevance is unknown.

### The mating inducer EroS_*Pv*_ enhances rosette development

We next investigated how single-celled *S. rosetta* in an SrEpac culture would respond to simultaneous exposure to EroS_*Pv*_ and RIF-OMVs. SrEpac cultures treated solely with RIF-OMVs for 0.5 hours, considerably less time than that required for rosette development, did not produce swarms and were indistinguishable from untreated SrEpac cultures (Fig. S1A-C’, Table 1; (18, 19)). Moreover, when SrEpac cultures were treated simultaneously with EroS_*Pv*_ and RIF-OMVs for 0.5 hours, the cells swarmed and the culture was indistinguishable from one treated with EroS_*Pv*_ alone (Fig. S1A, D-E’, Table 1). Therefore, RIF-OMVs do not appear to influence the swarm-inducing activity of EroS_*Pv*_ over time scales of 0.5 hours or less.

In contrast, when SrEpac cultures were co-treated with RIF-OMVs and EroS_*Pv*_ for 24 hours (long enough for rosettes to develop), the percentage of cells in rosettes increased markedly compared to cultures treated with RIF-OMVs alone (Fig. 2A, Table 1). Thus, EroS_*Pv*_ enhances the rosette-inducing activity of RIF-OMVs. The enhancing activity of EroS_*Pv*_ derived, in part, from the increased sensitivity of the culture to RIF-OMVs, allowing for rosette development at RIF-OMV concentrations that would otherwise fail to elicit rosette development. For example, at a nearly 10^−6^ dilution of RIF-OMVs, no rosettes were detected in the RIF-OMV alone condition, while 4.5±0.8% (mean ± S.D.) of the cells in cultures co-treated with EroS_*Pv*_ and RIF-OMVs were found in rosettes (see circle #1, Fig. 2A). In addition, when cells were exposed to saturating concentrations of RIF-OMVs (dilutions ≥3.7×10^−4^), co-treatment with EroS_*Pv*_ increased the percentage of cells in rosettes from a maximum of 83.6±6.8% (mean ± S.D.) in cultures that were treated with RIF-OMVs alone to 92.6±0.3% (mean ± S.D.) in cultures co-treated with RIF-OMVs and EroS_*Pv*_ (see circle #2, Fig. 2A).

**Figure 2:**
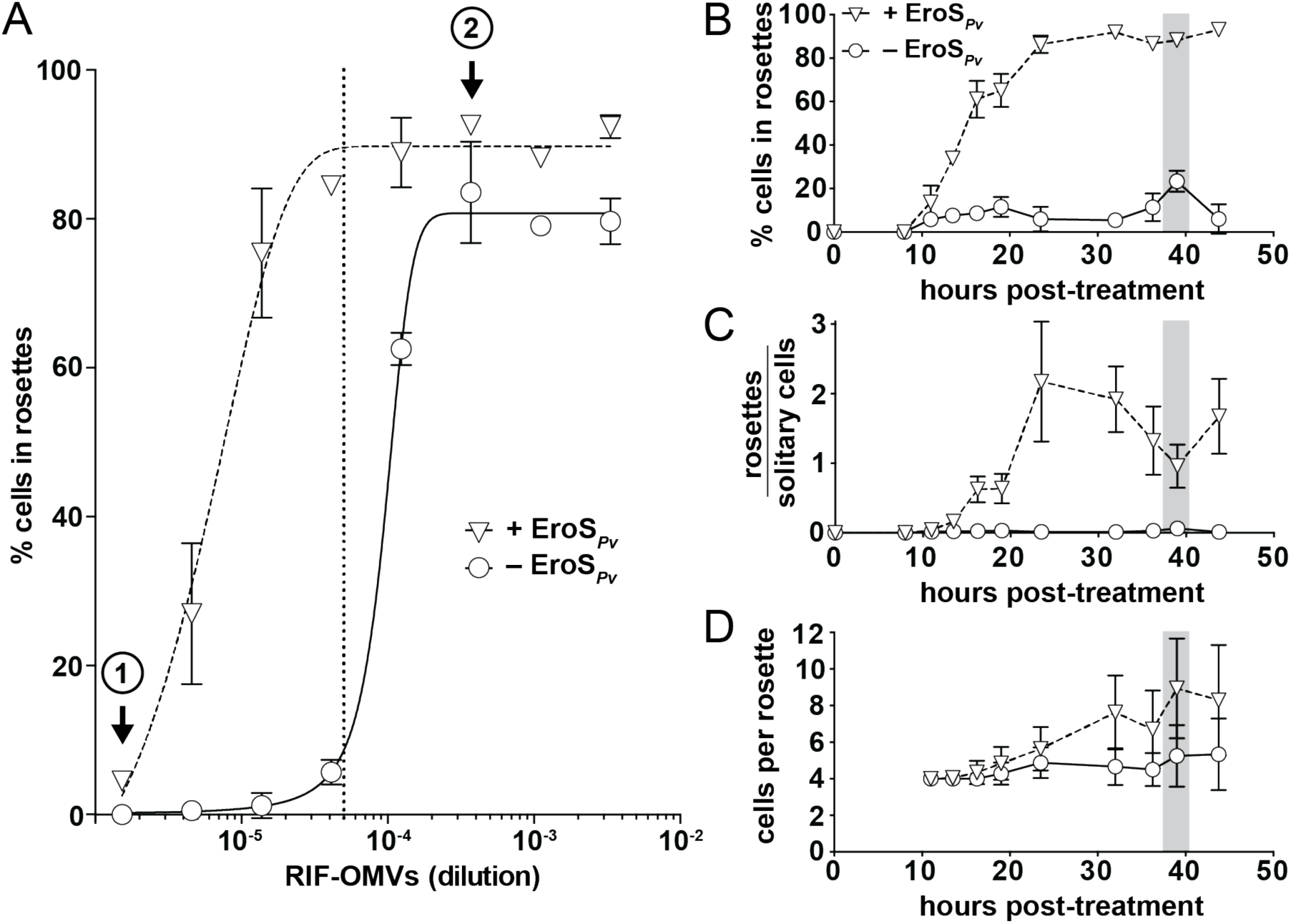
The mating inducer EroS_*Pv*_ enhances rosette development. (A) EroS_*Pv*_ enhances rosette induction by RIF-OMVs. Treatment of SrEpac with increasing concentrations of RIF-OMVs (circles) resulted in a concomitant increase in the percentage of cells in rosettes. Co-treatment of SrEpac with RIF-OMVs and 0.05 U/mL EroS_*Pv*_ (triangles) resulted in rosette development at concentrations of RIF-OMVs that did not otherwise induce rosettes (e.g. at (**1**)). EroS_*Pv*_ also increased the maximum percentage of cells in rosettes at saturating concentrations of RIF-OMVs (e.g. at (**2**)). The 1:20,000 dilution of RIF-OMVs used for the sensitized rosette induction assays in panels B-D is indicated with a vertical dotted line. Mean plotted ± S.D. (B) Co-treatment of SrEpac with EroS_*Pv*_ and RIF-OMVs leads to a dramatic increase in percentage of cells in rosettes throughout the course of rosette development relative to SrEpac treated only with RIF-OMVs. After 39 hours (shaded bar) of co-treatment with RIF-OMVs and EroS_*Pv*_ (triangles), 88.2±2.7% (mean ± S.D.) of *S. rosetta* cells were in rosettes, compared with 23.4±4.9% (mean ± S.D.) of cells treated with RIF-OMVs alone (circles). (C) EroS_*Pv*_ increased the ratio of rosettes to solitary cells in SrEpac cultures treated with RIF-OMVs. After 39 hours (shaded bar) of co-treatment with RIF-OMVs and EroS_*Pv*_ (triangles), the ratio of rosettes to solitary cells was 0.96±0.31 (mean ± S.D.), compared with 0.06±0.02 (mean ± S.D.) after treatment with RIF-OMVs alone (circles). (D) EroS_*Pv*_ increased the number of cells per rosette in RIF-OMV-treated SrEpac cultures. After 39 hours (shaded bar) of co-treatment with RIF-OMVs and EroS_*Pv*_ (triangles), there were 8.9±2.7 (mean ± S.D.) *S. rosetta* cells per rosette colony, compared with 5.3±1.7 (mean ± S.D.) cells per rosette colony after treatment with RIF-OMVs alone (circles).

Enhancement of rosette development by the mating factor EroS was unexpected, and we next sought to understand the phenomenon in greater detail. To that end, we optimized a method for reproducibly inducing rosette development at low levels. Treating SrEpac with a 1:20,000 dilution of RIF-OMVs drove only a small percentage of cells (1-20%) into rosettes (Fig. 2A, Fig. S2A) and thereafter formed the basis of a “sensitized rosette induction assay” in which we could quantify the influence of EroS_*Pv*_. Under the conditions of the sensitized rosette induction assay, we found that EroS_*Pv*_ enhanced rosette development in a concentration-dependent manner that saturated at 0.05 U/mL (Fig. S2B). Using this sensitized rosette induction assay across a time series, the rosette enhancing activity of EroS_*Pv*_ at the population level became more evident (Fig. S2C). For example, while treatment of SrEpac with 1:20,000 RIF-OMVs yielded only 23.4±4.9% (mean ± S.D.) of cells in rosettes at 39-hours post-treatment, co-treatment with 1:20,000 RIF-OMVs and 0.05 U/mL EroS_*Pv*_ yielded 88.2±2.7% (mean ± S.D.) of cells in rosettes (Fig. 2B).

These data demonstrated that co-treatment with EroS_*Pv*_ increases the percentage of cells in rosettes at a population level but did not reveal whether EroS_*Pv*_-mediated enhancement works by (1) increasing the overall number of rosettes, (2) increasing the average number of cells per rosette, or (3) both. To test whether co-treatment with EroS_*Pv*_ increased the number of rosettes formed, we induced SrEpac with either RIF-OMVs alone or RIF-OMVs + EroS_*Pv*_ and measured the ratio of rosette colonies to solitary cells. Co-treatment with RIF-OMVs and EroS_*Pv*_ in the sensitized rosette induction assay consistently increased the ratio of rosette colonies to solitary cells throughout the time series. For example, at 39 hours post-treatment, the ratio of rosettes to solitary cells after co-treatment with RIF-OMVs and EroS_*Pv*_ was 0.96±0.31 (mean ± S.D.), compared to 0.06±0.02 (mean ± S.D.) after treatment with RIF-OMVs alone (Fig. 2C). The ratio of rosettes to solitary cells eventually plateaued, likely due to both solitary cells and rosettes (which can divide by fission (20)) dividing at the same rate. To test whether rosette size is influenced by co-treatment with EroS_*Pv*_, we used the sensitized rosette induction assay to compare the number of cells per rosette in cultures treated with RIF-OMVs alone to those treated with RIF-OMVs and EroS_*Pv*_. Cultures co-treated with EroS_*Pv*_ formed larger rosettes (with 8.9±2.7 (mean ± S.D.) cells per rosette colony at 39-hours post-treatment) compared with those treated with RIF-OMVs alone (5.3±1.7 (mean ± S.D.) cells per rosette colony at the same time point) (Fig. 2D). Importantly, co-treatment with EroS did not affect the growth rate or cell density of cultures (Fig. S2D), indicating that the increase in cell number per rosette was not due to a difference in cell division rates. Therefore, at limiting concentrations of RIF-OMVs, EroS_*Pv*_ enhances the rosette-inducing activity of RIF-OMVs in at least two ways: at the population level, by increasing sensitivity to RIFs and the number of cells that initiate rosette development, and at the level of development, by increasing the maximal size of rosettes.

### Purified RIFs and EroS are sufficient for enhancement of rosette induction

Because *A. machipongonensis* OMVs contain a suite of proteins, sugars, the sulfonolipid RIFs, and diverse other lipids, we next explored whether RIFs are sufficient for EroS_*Pv*_-mediated enhancement of rosette development or whether the phenomenon requires a non-RIF. For example, certain lysophosphatidylethanolamines (LPEs), lipids found alongside RIFs in *A. machipongonensis* OMVs, synergize with RIFs and enhance rosette induction, in part by increasing the resistance of larger rosettes to shear forces (22).

To test whether EroS_*Pv*_ acts synergistically with RIFs or requires other components of RIF-OMVs, we compared rosette development in SrEpac cultures treated with high-performance liquid chromatography (HPLC)-purified RIFs (19,22) with that in cultures co-treated with HPLC-purified RIFs and EroS_*Pv*_. Co-treatment with EroS_*Pv*_ and purified RIFs caused a significant increase in the percentage of cells in rosettes compared to purified RIFs alone, indicating that the enhancement does not require other components of *A. machipongonensis* OMVs (Fig. 3A). Moreover, enhancement of rosette development was not restricted to *P. vulgaris EroS*. Co-treatment with purified *V. fischeri* EroS (EroS_*Vf*_) also significantly enhanced RIF-OMV-induced rosette development (Fig. 3B), revealing that the enhancing activity likely stems from the chondroitinase activity conserved between EroS_*Vf*_ and EroS_*Pv*_ rather than from a lineage-specific feature found only in EroS_*Pv*_. These findings show that simultaneous exposure to just two bacterial cues, RIFs and EroS, is sufficient to induce enhanced development of rosettes in *S. rosetta*.

**Figure 3:**
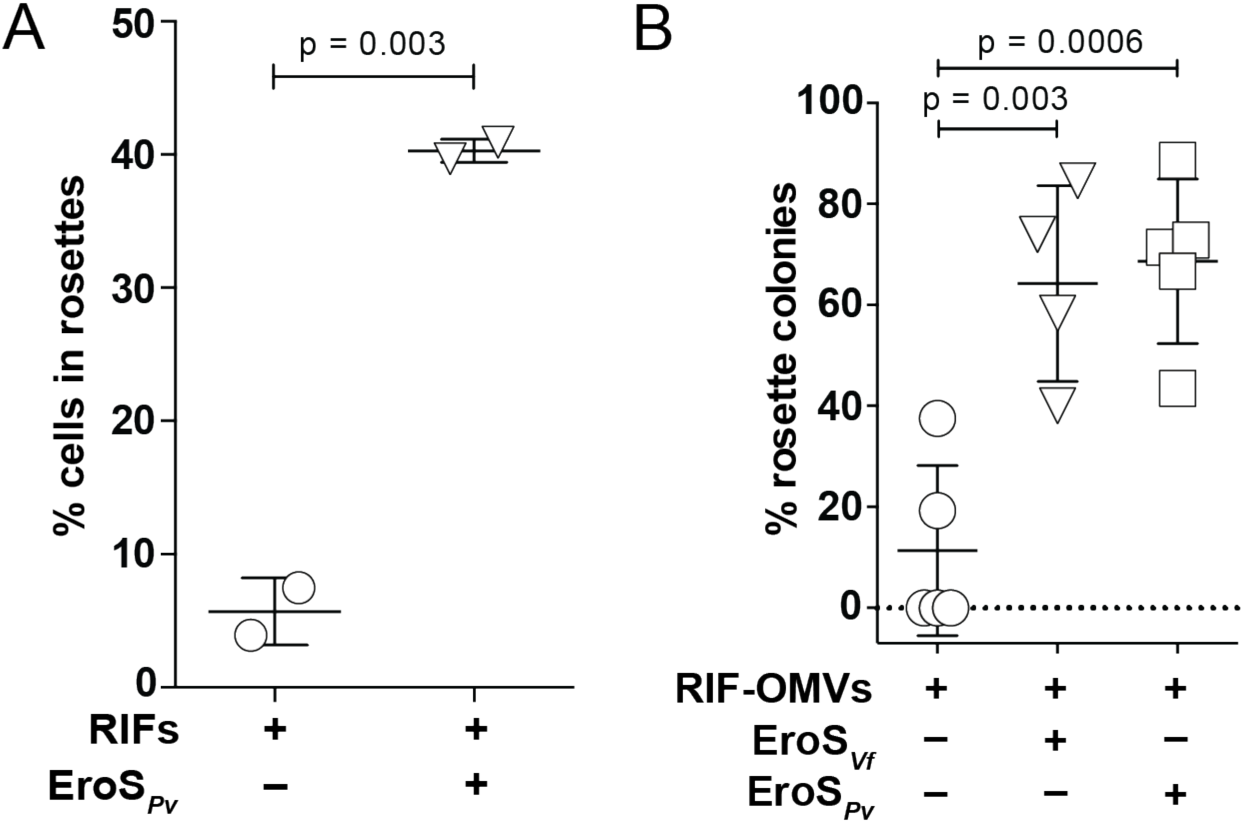
Purified RIFs and EroS are sufficient for enhancement of rosette induction. (A) Co-treatment of SrEpac with 10 µg/mL HPLC-purified RIFs and 0.05 U/mL EroS_*Pv*_ (triangles) resulted in an increase in the percentage of *S. rosetta* cells in rosettes compared to treatment with HPLC-purified RIFs alone (circles). Mean plotted ± S.D. (unpaired *t* test). (B) Co-treatment of SrEpac with a 1:20,000 dilution of RIF-OMVs and either 0.1% EroS from *V. fischeri* (EroS_*Vf*_), or 0.05 U/mL EroS from *P. vulgaris* (EroS_*Pv*_) resulted in an increase in the percentage of rosette colonies compared to treatment with RIF-OMVs alone. Mean plotted ± S.D. (unpaired *t* test).

## Discussion

We have shown here that the choanoflagellate *S. rosetta* can sense and respond to a mix of bacterial cues, each of which in isolation induces a seemingly disparate life history transition – mating or multicellularity. Together, these cues enhance multicellular development, increasing the number of cells in rosettes at a population level by increasing the proportion of rosettes to single cells and by increasing the number of cells per rosette (Fig. 2 and 4).

**Figure 4:**
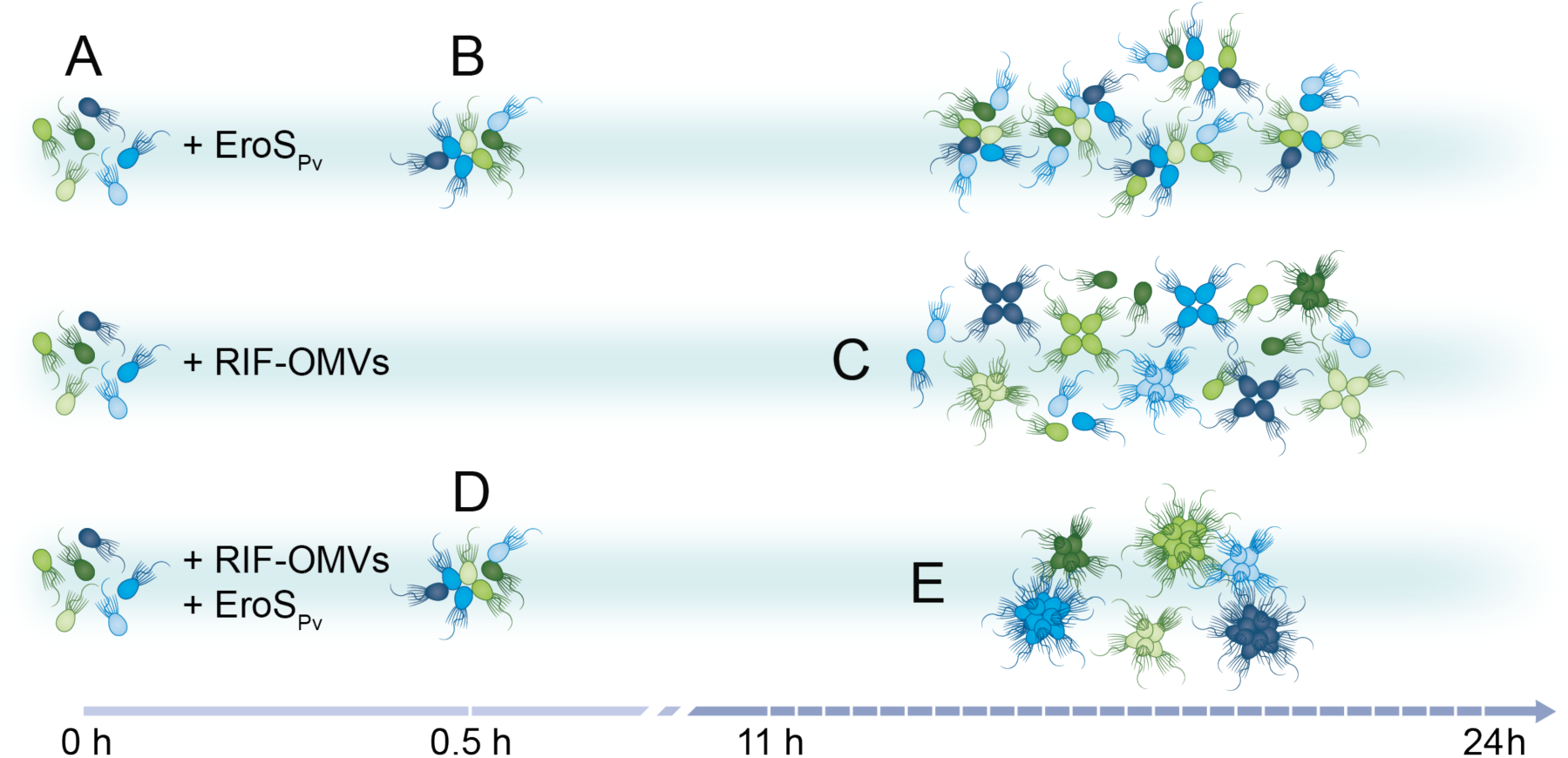
*S. rosetta* integration of bacterial cues. *S. rosetta* phenotypes induced over time by EroS_*Pv*_, RIF-OMVs, and the synergistic effect of both cues. (A) Untreated SrEpac proliferates as solitary cells. (B) Treatment with EroS_*Pv*_ induces swarming of unrelated cells within 0.5 hours. (C) Treatment with RIF-OMVs induces rosette development through cell division within 11-24 hours. (D) Co-treatment with RIF-OMVs and EroS_*Pv*_ for 0.5 hours results in swarming, showing that RIF-OMVs do not interfere with or enhance the activity of EroS_*Pv*_. (E) After 11-24 hours of co-treatment with RIF-OMVs and EroS_*Pv*_, rosettes develop and swarm. Compared to treatment with RIF-OMVs alone, co-treatment with RIF-OMVs and EroS_*Pv*_ induces the development of more rosettes and rosettes containing more cells.

The *S. rosetta* targets for EroS and the sulfonolipid RIFs are as-yet unknown (18, 19), making it challenging to infer the specific mechanisms by which EroS might enhance rosette development. One possibility is that EroS may modify chondroitin sulfate proteoglycans through its chondroitinase activity, thereby improving access of RIF receptors to RIFs, potentially explaining the increased sensitivity of EroS-treated *S. rosetta* to RIF-OMVs (Fig. 2A). This type of mechanism would resemble the regulation of vascular endothelial growth factor receptor 2 (VEGFR2), whose activity is inhibited by *N*-glycosylation; enzymatic digestion of glycans on VEGFR2 enhances its response to the VEGF ligand (25).

In addition to increasing the sensitivity of *S. rosetta* to RIF-OMVs, EroS treatment also resulted in rosettes that contained more cells (Fig. 2D). A link between rosette size and extracellular matrix (ECM) modification was previously reported for another colony-forming choanoflagellate, *Salpingoeca helianthica*, in which treatment with a bovine chondroitinase resulted in a significantly increased number of cells per rosette (26). Furthermore, chemical perturbations of the *S. rosetta* ECM and computational modeling have shown that the material properties of the ECM, such as stiffness and volume, exert a physical constraint on rosette volume and morphology (27). Thus, EroS digestion of chondroitin sulfate in the *S. rosetta* ECM may relax these constraints and allow for increased proliferation of cells within rosettes.

Might *S. rosetta* in nature actually encounter the disparate types of bacteria that induce multicellularity and mating? Rosette development can be induced by diverse genera of marine bacteria, ranging from *A. machipongonensis*, which was co-isolated with *S. rosetta*, to *Zobellia uliginosa*, a macroalgal commensal (19, 28, 29). Likewise, mating can be induced by diverse *Vibrio* species (18), which are widespread in marine environments (30, 31). Moreover, the bioactive molecules produced by *A. machipongonensis* and *V. fischeri* (sulfonolipid RIFs and EroS) are secreted and are potent at ecologically relevant concentrations (femtomolar to nanomolar) that are comparable to those of other soluble marine signaling molecules (2, 18, 19, 22). Taken together, the diversity and abundance of rosette-inducing and mating-inducing bacteria, and the potency of the molecules they produce, argue that RIFs and EroS could be simultaneously encountered by *S. rosetta* in nature. The synergy between these cues allows *S. rosetta* to sense and respond to significantly lower concentrations of rosette-inducing factors than it could otherwise (Fig. 2A), contributing to the plausibility that the enhanced rosette induction they elicit could be ecologically relevant.

Simple host-microbe interactions, in which a single bacterium elicits a clear phenotype from a eukaryotic host, have begun to reveal the molecular mechanisms by which bacteria influence the biology of eukaryotes. For example, *V. fischeri* colonizes and is sufficient to induce the development of the light organ in the bobtail squid, but this process only happens through the integration of multiple cues produced by *V. fischeri* – peptidoglycan and lipopolysaccharide (32). Likewise, we have previously shown that two types of molecules – sulfonolipid RIFs and specific LPEs – are necessary to recapitulate the rosette-inducing activity of live *A. machipongonensis* (22). Thus, interactions that are seemingly simple at the organismal level – one bacterium and one eukaryote – can require complex interactions at the molecular level.

Given the underlying molecular complexity of interactions involving only one bacterium and one eukaryote, interactions among larger numbers of species are, perhaps unsurprisingly, complex and can yield a variety of outcomes, including synergistic effects (12). For example, arbuscular mycorrhizal fungi and rhizobia bacteria individually confer beneficial effects on plants, and the simultaneous presence of both groups in a tripartite association enhances these effects, increasing plant biomass to a greater extent than each partner could alone (33). Synergistic effects have also been demonstrated in interactions among eukaryotes and multiple bacterial species, such as in polymicrobial infections. Direct interactions among pathogens in polymicrobial infections (through metabolite exchange, signaling molecules, or direct contact) can synergistically increase the disease burden for the host (such as by increasing antibiotic resistance or virulence factor expression) (34). Eukaryotic integration of bacterial cues has also been observed in the mammalian immune system, in which immune receptors such as Toll-like receptors, T cell receptors and co-receptors, each of which recognizes different bacterial ligands, synergize to enhance the response to multiple bacterial cues (35, 36).

Our finding that isolated cues from diverse environmental bacteria can synergize to enhance rosette development in *S. rosetta* (Fig. 3) demonstrates that this type of integration can occur at the level of the eukaryote, without requiring direct interactions among environmental bacteria. In the future, identifying the *S. rosetta* target(s) of RIF and EroS activity will likely provide detailed insights into the molecular mechanisms underlying EroS-mediated enhancement of rosette development. The experimental tractability of *S. rosetta* and its susceptibility to the influences of environmental bacteria render it an exciting system in which to investigate the mechanisms by which eukaryotes grapple with a noisy and information-rich bacterial world.

## Materials and Methods

### Choanoflagellate culturing conditions

Artificial seawater (ASW) was prepared by diluting 32.9 g Tropic Marin sea salts in 1L water for a salinity of 32-37 parts per thousand (24). Sea Water Complete media (SWC) was prepared by diluting 5 g/L peptone, 3 g/L yeast extract, and 3 mL/L glycerol in ASW (24). SrEpac (*Salpingoeca rosetta* co-cultured with the prey bacterium *Echinicola pacifica*, ATCC PRA-390; (24)) was cultured in 5% Sea Water Complete media (5% SWC vol/vol in ASW) at 22ºC. Cultures were passaged daily, 1 mL into 9 mL fresh media in 25cm^2^ cell culture flasks (Corning). Prior to rosette or swarm induction, cultures were diluted to 1×10^5^ choanoflagellate cells/mL in 5% SWC and 100 µL volumes were aliquoted into a 96-well plate (Corning).

### Preparation of *A. machipongonensis* conditioned media and isolation of RIF-OMVs

Outer membrane vesicles were isolated from *A. machipongonensis* as described in (22). Briefly, *A. machipongonensis* (ATCC BAA-2233, (28)) was grown in 500 mL 100% SWC, shaking at 30ºC for 48 hours. The bacteria were pelleted and the supernatant was filtered through a 0.2 µm filter to produce conditioned media. Conditioned media was then centrifuged at 36,000 × g for 3 hours at 4ºC (Type 45 Ti rotor, Beckman Coulter). OMV-containing pellets were resuspended in 2 mL ASW.

### HPLC purification of RIFs

RIFs were purified by HPLC as described in (22). Briefly, *A. machipongonensis* was grown in 20 L Marine Broth media (Carl Roth (CP.73): 40.1 g/L), shaking at 30ºC for 48 hours. The cells were harvested by centrifugation and extracted with CHCl_3_:MeOH (2:1, 4L). The organic extract was filtered and concentrated to give approximately 3g crude lipid extract. The crude extract was dissolved in 60% MeOH (+0.1% NH_4_OH) and fractionated using a C18-SPE (Solid Phase Extraction) using a 10% step-gradient of MeOH (60%-100% MeOH+0.1 NH_4_OH). The resulting SPE fractions were analyzed for sulfonolipid-specific signals using LC-MS and ^1^H-NMR. The fraction containing RIF-mix (RIF-1 and RIF-2) eluted with 90% MeOH (+0.1% NH_4_OH) during the SPE purification.

### Rosette induction

Unless otherwise noted, SrEpac cultures were treated with a 1:1,000 dilution of RIF-OMVs and incubated for 24 hours before imaging or counting. To induce a low level of rosette development in the sensitized rosette induction assay (Fig. 2B-D, Fig. 3B, Fig. S2B-D), SrEpac cultures were treated with a 1:20,000 dilution of RIF-OMVs. HPLC-purified RIFs were resuspended in DMSO and added at 10 µg/mL (Fig. 3A).

### Swarm induction

Unless otherwise noted, cultures were treated with 0.05 U/mL chondroitinase ABC from *P. vulgaris* (Sigma), referred to as “EroS_*Pv*_”. EroS from *V. fischeri* (EroS_*Vf*_; Fig. 3B) was purified as described in (18). Briefly, *V. fischeri* ES114 (ATCC 700601) was grown in 8 L 100% SWC, shaking at 20ºC for 30 hours. The bacteria were pelleted and the supernatant was filtered through a 0.2 µm filter, concentrated to 120 mL using a using a tangential flow filtration device with a 30 kDa Centramate filter (Pall #OS030T12), then ammonium sulfate precipitated and further separated by size exclusion chromatography. EroS_*Vf*_ was added to SrEpac cultures at a final dilution of 0.1%.

### Rosette quantification

To quantify the percentage of cells in rosettes, cultures were fixed with 1% formaldehyde, vortexed, mounted on a Bright-Line hemacytometer (Hausser Scientific), and counted on a Leica DMI6000B inverted compound microscope. Rosettes were defined as groups of four or more cells, and were distinguished from swarms based on their resistance to mechanical shear and their stereotypical orientation with their basal poles pointed inwards and their flagella out (20, 23). The numbers of solitary cells and rosettes, as well as the number of cells in each rosette, were counted until at least 200 cells were scored (per biological replicate).

### Swarm quantification

Cell cluster areas were quantified as described in (18). Briefly, samples were imaged in 96-well glass-bottomed plates (Ibidi 89621) at 10× magnification using transmitted light (bright field) on a Zeiss Axio Observer.Z1/7 Widefield microscope with a Hammatsu Orca-Flash 4.0 LT CMOS Digital Camera. Images were processed and analyzed in ImageJ as follows: ‘Smooth’ to reduce bacterial background, ‘Find Edges’ to further highlight choanoflagellate cells, ‘Make Binary’ to convert to black and white, ‘Close-’ to fill in small holes, and ‘Analyze Particles’ to calculate the area of each cell cluster. Particles smaller than 10 µm^2^ were removed to reduce background bacterial signal.

## Acknowledgements

This material is based upon work originally supported by the National Institutes of Health (R01-GM099533 to N.K.). We thank members of the King lab for critical feedback, especially David Booth, Thibaut Brunet and Ben Larson. We thank Christine Beemelmanns and Chia-Chi Peng for providing HPLC-purified RIFs, Zoe Vernon and Olivia Angiuli for statistics consultation, and Debbie Maizels for help with illustrations in Fig. 4.

## SUPPLEMENTAL MATERIAL

**Figure S1:**
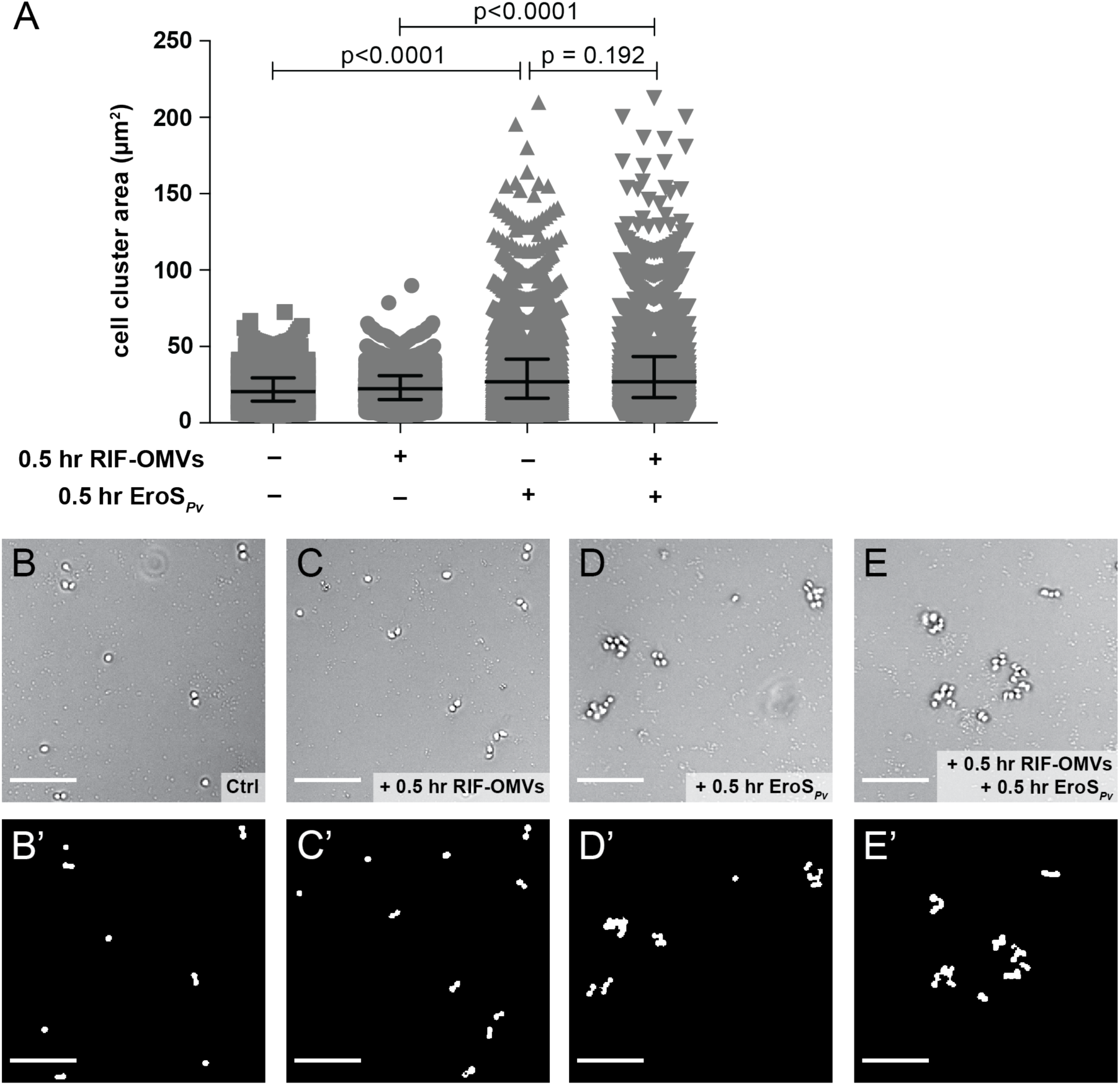
RIF-OMVs have no effect on EroS_*Pv*_-induced swarming. (A) Solitary cells from SrEpac co-treated with RIF-OMVs and EroS_*Pv*_ formed swarms, quantifiable by an increase in cell cluster area (median = 27.0 µm^2^, interquartile range = 16.5-43.5 µm^2^) compared to cells treated with RIF-OMVs alone (median = 22.4 µm^2^, interquartile range = 15.2-30.8 µm^2^). There was no significant difference in swarm size between cells co-treated with RIF-OMVs and EroS_*Pv*_ and cells treated with EroS_*Pv*_ alone (median = 27.0 µm^2^, interquartile range = 16.0-41.8 µm^2^) (Kolmogorov-Smirnov test). A minimum of 2730 cell cluster areas were plotted for each condition. (B-E’) Sample images used for quantification in (A). Following the approach of (18), raw images in (B-E) were converted to binary images (B’-E’) to measure cell cluster size (Materials and Methods). (B) *S. rosetta* cells from untreated SrEpac remained solitary. (C) *S. rosetta* cells from SrEpac treated with RIF-OMVs for 0.5 hours remained solitary. (D) *S. rosetta* cells from SrEpac treated with EroS_*Pv*_ for 0.5 hours formed visible swarms. (E) *S. rosetta* cells from SrEpac co-treated with RIF-OMVs and EroS_*Pv*_ for 0.5 hours formed visible swarms.

**Figure S2:**
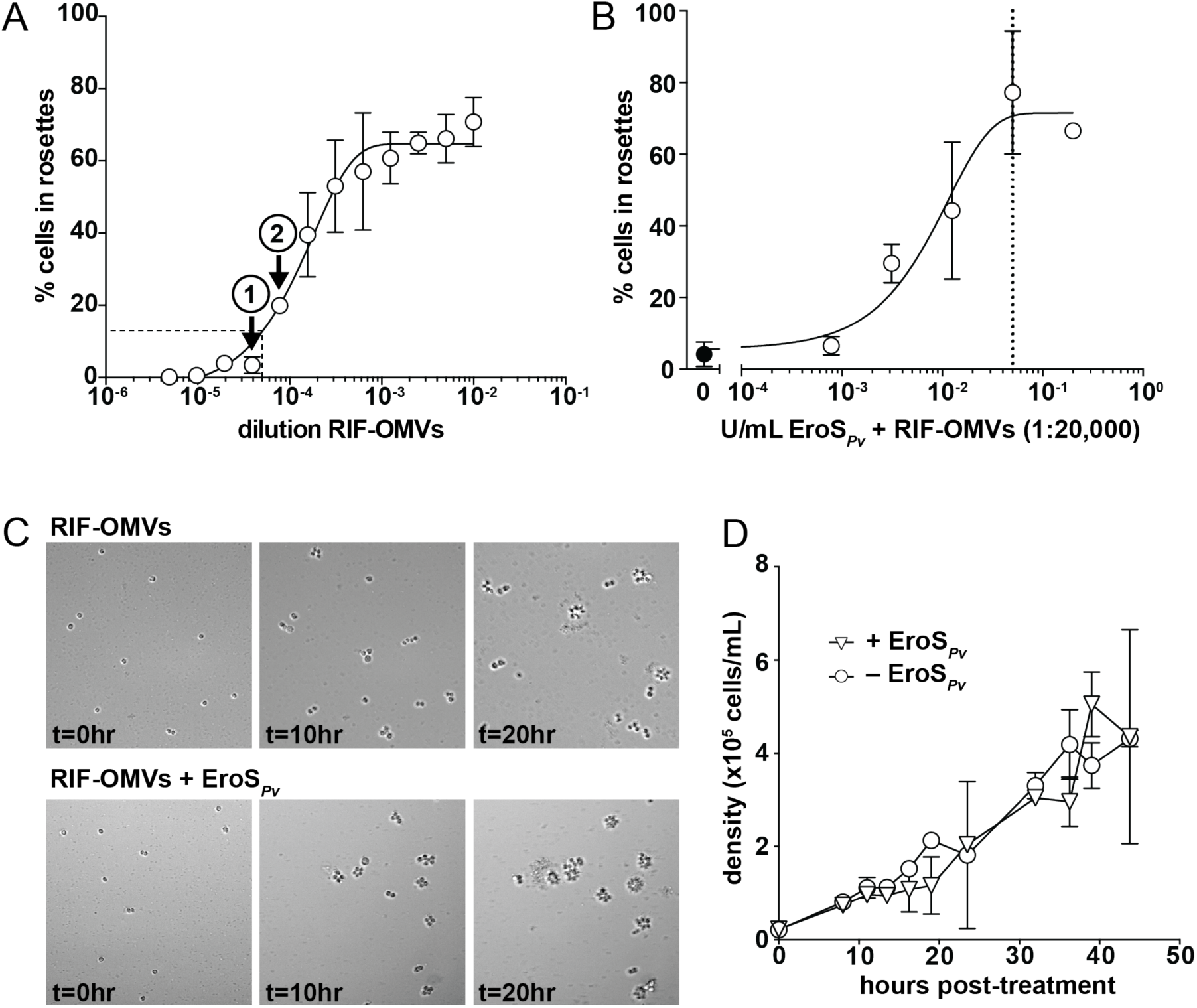
EroS_*Pv*_ enhances rosette development, but not cell proliferation, in a sensitized rosette induction assay. (A) Serial dilution of RIF-OMVs can be used to induce a low percentage of cells in rosettes. SrEpac treated with a 1:25,600 dilution of RIF-OMVs resulted in 3.4±2.3 (mean ± S.D.) *S. rosetta* cells in rosettes (arrow marked (**1**)), while a 1:12,800 dilution of RIF-OMVs resulted in 19.9±1.7 (mean ± S.D.) *S. rosetta* cells in rosettes (arrow marked (**2**)). An intermediate dilution of 1:20,000 was used for the sensitized rosette induction assay (dashed lines). (B) Rosette-enhancing activity correlated with EroS_*Pv*_ concentration. SrEpac treated with a 1:20,000 dilution of RIF-OMVs (black circle) contained more *S. rosetta* cells in rosettes upon the addition of increasing concentrations of EroS_*Pv*_ (white circles). Dotted line indicates concentration of EroS_*Pv*_ (0.05 U/mL) used for subsequent assays. Mean plotted ± S.D. (C) Time-lapse imaging showed an increase in both the number of rosettes and the number of cells per rosette after co-treatment with a 1:20,000 dilution of RIF-OMVs and 0.05 U/mL EroS_*Pv*_ (bottom) compared to RIF-OMVs alone (top). (D) *S. rosetta* cells treated with RIF-OMVs (circles) or co-treated with RIF-OMVs and EroS_*Pv*_ (triangles) grew at the same rate. Mean density plotted ± S.D.

